# In silico optimization of RNA-protein interactions for CRISPR-Cas13-based antimicrobials

**DOI:** 10.1101/2022.06.21.496952

**Authors:** Ho-min Park, Yunseol Park, Urta Berani, Eunkyu Bang, Joris Vankerschaver, Arnout Van Messem, Wesley De Neve, Hyunjin Shim

**Affiliations:** Center for Biosystems and Biotech Data Science, Ghent University Global Campus, Incheon, South Korea; Department of Electronics and Information Systems, Ghent University, Ghent, Belgium; Department of Applied Mathematics, Computer Science and Statistics, Ghent University, Ghent, Belgium; Department of Mathematics, University of Liège, Liège, Belgium

**Keywords:** CRISPR-based antimicrobials, RNA-protein interaction, RNA secondary structure, RNA tertiary structure, in silico docking, drug design, structural biology

## Abstract

RNA-protein interactions are crucial for diverse biological processes. In prokaryotes, RNA-protein interactions enable adaptive immunity through CRISPR-Cas systems. These defense systems utilize CRISPR RNA (crRNA) templates acquired from past infections to destroy foreign genetic elements through crRNA-mediated nuclease activities of Cas proteins. Thanks to the programmability and specificity of CRISPR-Cas systems, CRISPR-based antimicrobials have the potential to be repurposed as new types of antibiotics. Unlike traditional antibiotics, these CRISPR-based antimicrobials can be designed to target specific bacteria and minimize detrimental effects on the human microbiome during antibacterial therapy. Here, we explore the potential of CRISPR-based antimicrobials by optimizing the RNA-protein interactions of crRNAs and Cas13 proteins. CRISPR-Cas13 systems are unique as they degrade specific foreign RNAs using the crRNA template, which leads to non-specific RNase activities and cell cycle arrest. We show that a high proportion of the Cas13 systems have no colocalized CRISPR arrays, and the lack of direct association between crRNAs and Cas proteins may result in suboptimal RNA-protein interactions in the current tools. Here, we investigate the RNA-protein interactions of the Cas13-based systems by curating the validation dataset of Cas13 protein and CRISPR repeat pairs that are experimentally validated to interact, and the candidate dataset of CRISPR repeats that reside on the same genome as the currently known Cas13 proteins. To find optimal CRISPR-Cas13 interactions, we first validate the 3-D structure prediction of crRNAs based on their experimental structures. Next, we test a number of RNA-protein interaction programs to optimize the in silico docking of crRNAs with the Cas13 proteins. From this optimized pipeline, we find a number of candidate crRNAs that have comparable or better in silico docking with the Cas13 proteins of the current tools. This study fully automatizes the in silico optimization of RNA-protein interactions as an efficient preliminary step for designing effective CRISPR-Cas13-based antimicrobials.

## Introduction

The central dogma of molecular biology attributes the main role of RNA as the intermediate messenger between DNA and protein [1]. Recent studies reveal that RNA is involved in diverse cellular processes such as regulatory activities of gene expression, catalytic activities of various substrates, and molecular chaperoning and scaffolding [2]. The ability of RNAs to interact with RNA-binding proteins (RBPs), which rely on both RNA sequence and structure, has been studied in a number of RNA-protein complexes, including ribosomal RNA complexes [3]. In prokaryotes, RNA-protein interactions play a vital role in the highly intricate process of adaptive immunity against foreign genetic elements through Clustered Regularly Interspaced Short Palindromic Repeats (CRISPR) and CRISPR-associated system (Cas) proteins [4,5]. Prokaryotic genomes with CRISPR-Cas systems have the ability to store sequence information of previous infections in their CRISPR arrays. A complex of Cas proteins uses this sequence information as a genetic template to find and neutralize invaders of the same sequence. To achieve such specificity, the interaction between a complex of Cas proteins with nuclease activities and a CRISPR RNA (crRNA) with the protein-binding component (CRISPR repeat) is essential for this RNA-mediated adaptive immunity. The redesign of the protein-binding component of crRNAs associated with the Cas9 protein (trans-activating RNA) led to an efficient genome-editing tool in diverse eukaryotic cells [6–8].

Thanks to the interest in genome-editing applications, there was an active discovery of new CRISPR-Cas systems based on Cas proteins, which revealed the immense diversity of CRISPR-Cas systems in nature [9–11]. Recently, CRISPR-Cas systems are being repurposed as antibiotic tools against multidrug-resistant bacteria due to their programmability and specificity [12–15]. The uncontrolled spread of antimicrobial resistance (AMR) against traditional antibiotics of small molecules has become a global health issue [16], and we are in urgent need of novel antibiotics to combat multidrug-resistant bacteria. Some novel strategies are bacteriophage-derived, such as phage therapy that has been used successfully to treat multidrug-resistant infections as breakthrough therapy [17–19]. Several phage-derived endolysins are in clinical development for their antimicrobial activities to lyse the peptidoglycan layer of Gram-positive bacteria [20,21]. Another promising strategy is to use bacterial defence systems such as CRISPR-Cas systems against themselves by reprogramming CRISPR templates to target AMR genes in the chromosome or on plasmids [12,13,22]. For this purpose, Cas9 proteins have been explored extensively, but the double-stranded breaks in DNA resulting from their nuclease activities leave the blunt-ends susceptible to DNA repair pathways in bacteria [23,24].

Recently, Class 2 CRISPR-Cas systems of type VI are being investigated as promising antimicrobial tools, whose activity is characterized by RNA-guided single-stranded RNA (ssRNA) cleavage [25,26]. These systems encompass a single-effector Cas13 protein consisting of two Higher Eukaryotes and Prokaryotes Nucleotide-binding (HEPN) domains with ribonuclease activity. Cas13 proteins bind and cleave specific RNAs, which sequentially activate nonspecific RNase activities by changing their structural conformation [26,27]. Such promiscuous RNA cleavage is effective in restricting bacteria growth by degrading bacterial transcript RNAs. As the effects of CRISPR-Cas13 systems cannot be repaired like those of CRISPR-Cas9 systems in prokaryotes, they are one of the most promising antimicrobial tools to resensitize and neutralize multidrug-resistant bacteria. A recent experimental study demonstrated that CRISPR-Cas13a systems could be designed to trigger such activities in a sequence-specific manner that led to successful bacteria growth arrest [28]. Currently, CRISPR-Cas13 systems are divided into five subtypes (VI-A, VI-B1, VI-B2, VI-C, VI-D), depending on the Cas13 protein and its accessory proteins. However, the architecture of CRISPR-Cas13 systems is highly variable [11].

In this study, we observe that many CRISPR-Cas13 systems have no adjacent CRISPR arrays, particularly the Cas13a systems. This genomic architecture implies that CRISPR-Cas13 systems often share CRISPR arrays with other CRISPR-Cas systems in the genome, and some structural studies used synthetic constructs due to the lack of clear association between the Cas13 protein and the crRNA [29]. For effective CRISPR-based antimicrobials, RNA-protein interactions between crRNAs and Cas proteins should be optimal. Previously, it was shown that some off-target effects of genome-editing tools in cells occur when there is a competition between several crRNAs to bind the Cas protein [30]. As CRISPR-based antimicrobials have to operate in bacterial cells which often have several CRISPR arrays, it is imperative that these antibiotic tools have the optimal affinity between the target crRNA and the Cas13 protein to prevent competition with endogenous crRNAs. Here, we optimize RNA-protein interactions by curating crRNA datasets from CRISPR-Cas13 genomes and predicting 3-D crRNA structures for in silico docking with the Cas13 protein. First, we compare the accuracy performance of several RNA secondary and tertiary structure prediction programs using the experimental data of crRNA and Cas13 structures. Next, we validate in silico docking of crRNAs on the Cas13 protein of interest using the experimental structures and the predicted structures of crRNAs. This comparison study optimizes the computational pipeline required for in silico docking experiments to assess RNA-protein interactions. Finally, we conduct in silico docking of candidate crRNAs on the Cas13 protein of interest to compare with the experimental data of the crRNA-Cas13 complexes. This candidate study contributes to the investigation of effective CRISPR-based antimicrobials by generating a list of candidate crRNAs that dock optimally with the Cas13 proteins. Furthermore, we aim to provide an optimized and automatized computational pipeline for in silico docking experiments to model the receptor-ligand binding of experimental or predicted structures. Such in silico studies offer an efficient preliminary step to scan for candidate crRNAs predicted to bind optimally with the Cas13 proteins to be validated and optimized further with in vitro and in vivo studies.

## Materials and Methods

### Genomic architecture of CRISPR-Cas13 systems

Two distinct components are necessary for a functional CRISPR-Cas system: a CRISPR array and a cluster of *cas* genes arranged in one or more operons (Figure 1). The CRISPR array consists of almost identical and mostly palindromic repeats, which are separated by unique spacers that contain foreign DNA from past infections. The *cas* genes are divided into four functional modules: the adaptation module of spacer acquisition, the expression module of pre-crRNA processing, the interference module of target recognition, binding and cleavage, and the signal transduction module of CRISPR-linked accessory genes. Currently, CRISPR-Cas systems are assigned to a class and type based on the composition of functional modules. In Class 1 (types I, III and IV), the effector module (part of the interference module) consists of multiple Cas proteins, whereas in Class 2 (types II, V and VI), a single and large Cas protein is responsible for the effector module [11].

**Figure 1:**
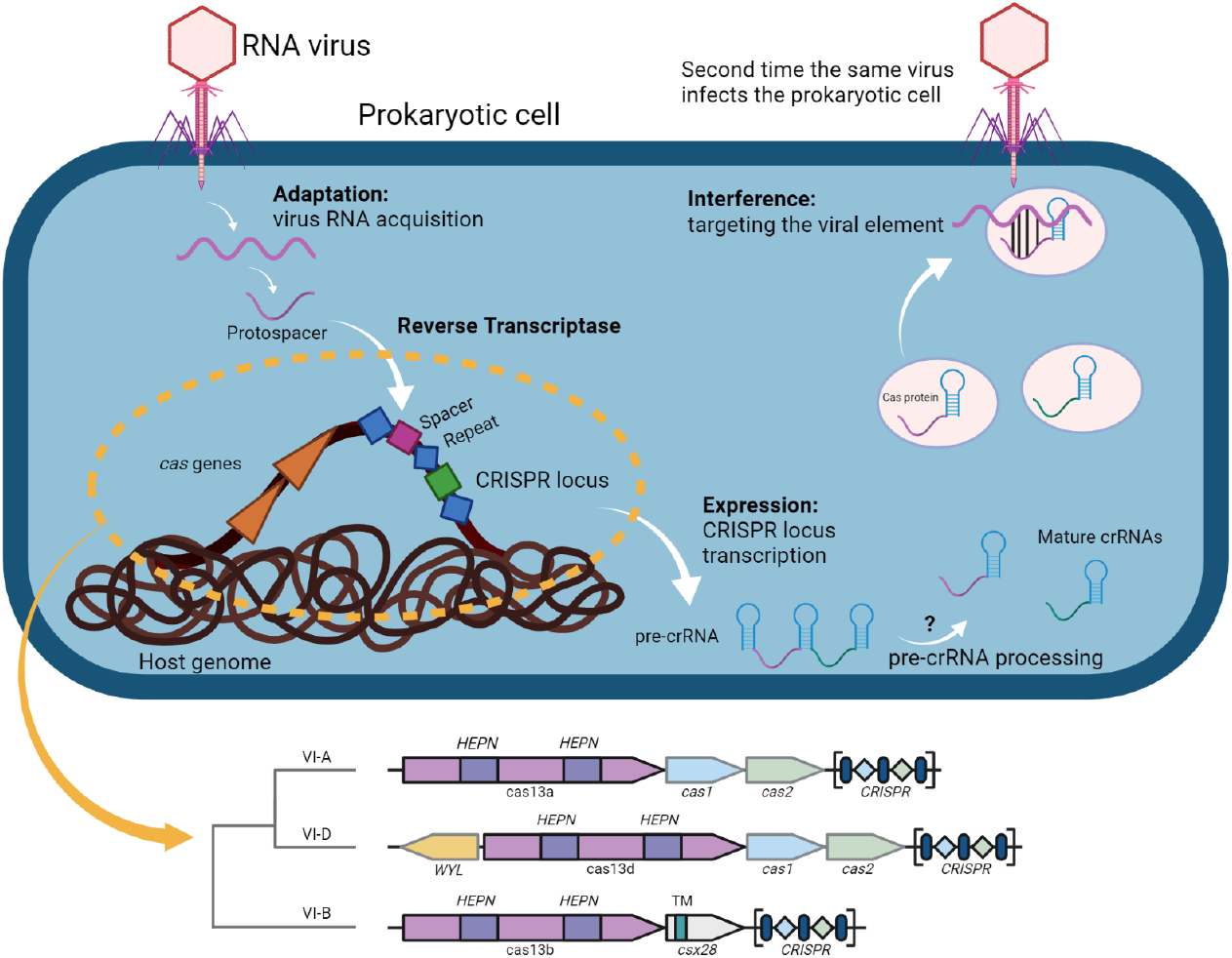
Architecture and mechanism of CRISPR-Cas13 systems. Three main stages constitute the CRISPR-Cas13 immune response: adaptation, expression and interference. During the adaptation stage, a complex of Cas proteins binds the invading genome, which is shown as an RNA virus. The bound part of the target RNA is cleaved out and is inserted into the CRISPR array of the prokaryotic genome as a new spacer through a reverse transcriptase. The expression stage involves the transcription of the CRISPR array as a large, single transcript and this pre-crRNA is processed into a mature crRNA containing a target spacer and a flanking repeat. The mechanisms and components involved in the pre-crRNA processing of CRISPR-Cas13 systems have not been experimentally resolved yet. At the last stage of the immune response, the interference stage utilizes the crRNA as a guide to recognize invading genomes based on sequence complementarity, recruiting the complex of Cas proteins. The Cas13a/b/d proteins have two higher eukaryotes and prokaryotes nucleotide-binding (HEPN) domains of RNase activity, which cleave the target sequence and inactivate the RNA virus.

CRISPR-Cas13 systems belong to Class 2 and type VI with several subtypes of effector proteins, including Cas13a, Cas13b, and Cas13d [26]. The Cas13 effector proteins contain two higher eukaryotes and prokaryotes nucleotide-binding (HEPN) domains that confer RNase activity (Figure 1). The Cas13 protein complexes with the crRNA via the CRISPR repeat sequence of ∼30 nucleotides, and the CRISPR spacer encodes a sequence that is complementary to the target sequence. In Class 2, CRISPR-Cas systems mostly involve Cas1, Cas2 and Cas4 for adaptation, but the Cas13b subtype lacks the adaptation module (Figure 1). Interestingly, the Cas13 family often has no colocalized CRISPR arrays within ±10,000 base pairs (Table S1). Compared to the Cas9 family with a high occurrence of colocalized CRISPR arrays (80%), the Cas13 family has lower occurrences of colocalized CRISPR arrays, particularly the subtypes Cas13a (19%) and Cas13d (0%).

### Data curation of Cas13 proteins and CRISPR repeats

To conduct in silico docking experiments, the first step was to curate a validation dataset of Cas13 proteins and associated CRISPR repeats that could be used for performance evaluation (Figure 2). An experimental dataset of the Cas13 family was retrieved from the Protein Data Bank (PDB), whose 3-D structure has been resolved by experimental techniques such as X-ray crystallography, NMR spectroscopy, or cryo-electron microscopy (Table S2). The repeat sequences of the crRNAs in the CRISPR-Cas complex were retrieved from the PDB in the FASTA format (Table S3). The CRISPR repeat sequences of all Cas13a-associated complexes were later shortened to match the visible part in the PDB, as their 3-D structures were only partially modelled. Some CRISPR-Cas13 systems contained multimeric proteins, and the crRNA chains (5W1I, 7OS0, 6IV8) were considered separately for the validation studies.

**Figure 2:**
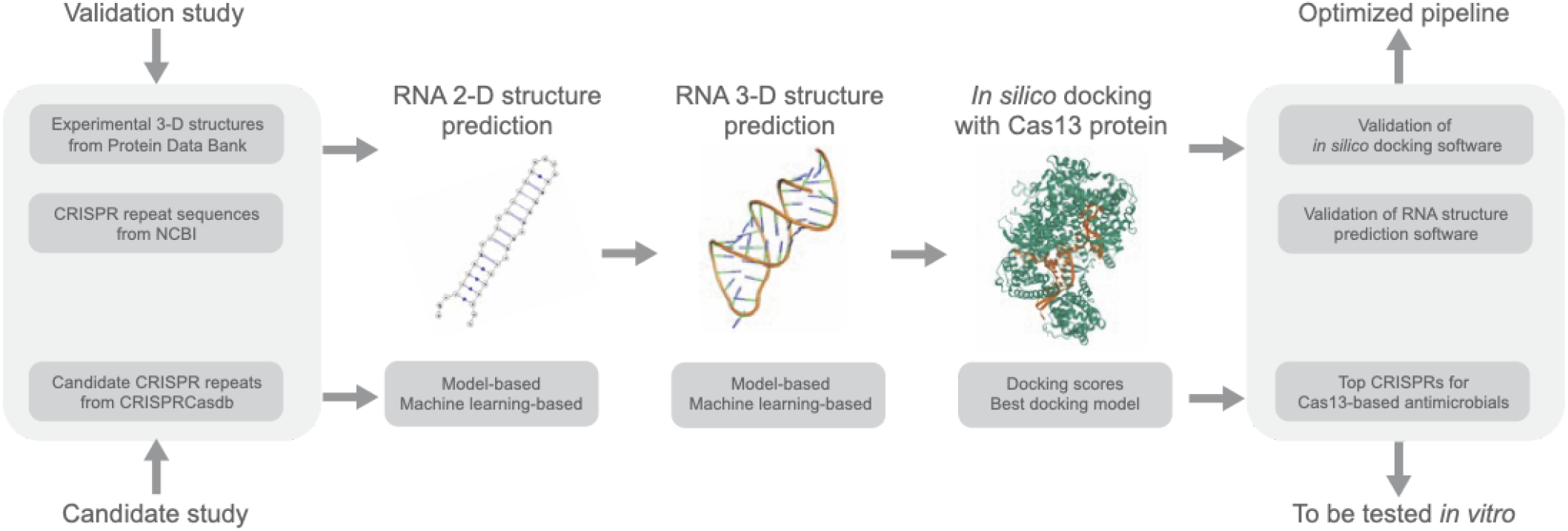
In silico docking of crRNAs with Cas13 proteins to assess the RNA-protein interactions. This study is divided into two parts: the validation study to optimize the in silico docking of crRNAs and Cas13 proteins, and the candidate study to apply the optimized pipeline of in silico docking to test a list of candidate crRNAs on each Cas13 protein for RNA-protein interactions, as a preliminary step prior to experimental validation.

Another dataset of candidate CRISPR repeats was curated to identify crRNAs that interact optimally with the Cas13 proteins. As shown previously, the Cas13 family often has no colocalized CRISPR arrays, which makes the association between the Cas13 proteins and the crRNAs difficult to determine (Figure S1). Thus, this candidate dataset is a collection of CRISPR repeat sequences from the CRISPR arrays that are within ±10,000 base pairs of the Cas13 proteins by querying the prokaryotic genomes from the CRISPRCasdb [31]. As these crRNAs have no experimental structures, the first step to in silico docking experiments is to predict the RNA structures of these candidate crRNAs.

### 3-D structure prediction of CRISPR repeats

As the RNA 2-D structure prediction precedes the 3-D structure prediction, the 3-D structure prediction programs base their prediction on the 2-D structure (Figure S2). With a selection of eight RNA 2-D structure prediction programs and two RNA 3-D structure prediction programs, 16 different combinations, including both machine learning-based and model-based methods, were evaluated using the validation dataset (Tables S4-S6). The 2-D structure of each CRISPR repeat sequence was predicted as the dot-bracket notation through the web servers of all the RNA 2-D structure prediction programs with the default parameters (Table S4).

RNAComposer receives genetic sequence and 2-D structure as input to predict the 3-D structure of an RNA molecule. If only the genetic sequence is given as an input, RNAComposer creates the 2-D structure itself through the built-in algorithm [32,33]. For Rosetta, the fragment assembly of RNA with full-atom refinement (FARFAR2) protocol was applied followingly [34]. The CRISPR repeat sequences of the validation dataset and the dot-bracket notation of the predicted 2-D structure were given as an input. Subsequently, the predicted RNA 3-D models were subjected to minimization in an all-atom scoring function used by the FARFAR2 protocol. The FARFAR2 protocol selected the best model in terms of minimum free energy (MFE) and root-mean-square deviation (RMSD) from the ensemble of predicted 3-D structure models. First, the models were sorted by total energy, and the top 500 models were selected. Then, these top 500 models were sorted by the RMSD value to extract the best 3-D model based on the RMSD and MFE values. The final output of the validation dataset resulted in 256 predicted crRNA 3-D structures in PDB files, given 16 crRNA sequences and 16 different RNA programs.

### Evaluation of the 3-D structure prediction of CRISPR repeats

To assess the performance of each program combination, the predicted 3-D structures of the crRNAs in the validation dataset were compared with the experimental 3-D structures (referred to as ‘ground truth’ or ‘GT’). The RMSD measuring the average distance between the atoms of superimposed structures was used as the performance metric. The RMSD values were obtained using three different evaluation programs: the PyMOL align function, the PyMOL super function and the secondary structure-based tertiary structure similarity algorithm (SETTER) [35,36]. The PyMOL align function superimposes 3-D structures based on sequence, while the PyMOL super function and the SETTER superimpose 3-D structures based on structure.

Each of the 256 predicted 3-D structures was superimposed with the corresponding GT structure, and this performance analysis was conducted with three replicates to account for the stochasticity of each evaluation program. The mean and standard deviation (SD) of the RMSD values across all predicted structures and all replicate runs were calculated as summary statistics. By keeping the RNA 3-D structure prediction program constant and changing the 2-D program, the prediction accuracy of each combination could be assessed. The best-performing combination with the lowest mean RMSD and SD value was chosen for further in silico docking experiments with the candidate dataset.

### In silico docking of experimentally validated crRNAs with Cas13 proteins

The validation dataset contained three subtypes of Cas13: Cas13a, Cas13b, and Cas13d. These experimentally validated structures from the PDB contained CRISPR-Cas complexes with both the Cas protein and crRNA. The PDB files of the validation dataset were first cleaned to remove all unwanted residues such as water and magnesium molecules in PyMOL. For the PDB files containing dimeric Cas proteins and two crRNAs (5W1I, 6IV8, 7OS0), the two chains and RNAs were separated and named with their chain ID (5W1I_AB, 5W1I_CD; 6IV8_AB, 6IV8_CD; 7OS0_AF, 7OS0_CD). Each of the cleaned PDB files were separated into the receptor (Cas protein) and ligand (crRNA).

Next, the receptor and ligand pairs were randomly rotated to ensure that the docking is not influenced by their initial coordinates. For example, PyDockDNA without random rotation led to perfect docking of the ligand and the receptor. Each receptor and ligand pair was rotated separately in 3-D around the center of mass using random angles between 40-320°, to ensure sufficient rotation from the initial coordinates. Three different angles were chosen for the Cas proteins and for each crRNA.

To optimize in silico docking using the validation dataset, five in silico docking programs were considered: HADDOCK [37,38], HDOCK [39], PyDockDNA [40], RNP-denovo [41], and Swarmdock [42]. However, two of these were eliminated in the preliminary steps due to software incompatibility and thus, only three programs (HADDOCK, HDOCK, and PyDockDNA) were considered for the in silico docking validation study (Table S7). We automatized the use of the web servers for in silico docking experiments. HADDOCK was run with the parameters recommended for RNA-protein docking. Some experimental structures with intrinsically disordered proteins were processed prior to using HADDOCK, as the dynamic conformations of the proteins intervened with in silico docking [43]. All other parameters were kept as default, except for the parameter defining randomly ambiguous interaction restraints from accessible residues, which allows docking without specifying the binding sites. PyDockDNA was run with the default parameters and the PyDock scoring function was used. HDOCK was run with the template-free option, and all other parameters were kept as default. For evaluation of the results, only the best docking model, as given by the internal docking score based on a relative ranking of different binding models for the receptor-ligand pair, was considered in the validation study.

In the following steps of the validation study, the predicted crRNA structures were used as ligands. These RNA 3-D structures were predicted with the shortened sequences containing only the CRISPR repeat. The RNA 3-D structures were generated from all combinations of the 2-D and 3-D structure prediction programs (Table S6). The subsequent in silico docking of the receptor (Cas13a proteins) and the ligand (predicted crRNA structures) was conducted template-free and template-based runs of HDOCK.

We used the interface RMSD (iRMSD) to assess the performance of each in silico docking experiment, which is one of the docking scores used in Critical Assessment of PRediction of Interactions (CAPRI) experiments [44,45] that calculates the docking distance overlap between the GT ligand and the model ligand at the interface of all atoms within a distance of 8 Å from the point where the receptor and ligand meet (see Figure S3 for details). The CAPRI-based performance calculation was adapted for RNA-protein interactions by retaining alpha carbon (Cα) and phosphate as the backbone of the protein receptor and the RNA ligand, respectively [46]. Finally, we used DockQ [47] for calculating iRMSD values by setting the backbone atom as Cα and phosphate.

### In silico docking of candidate crRNAs with Cas13 proteins

The candidate crRNAs structures of Cas13a, Cas13b, and Cas13d were predicted with the two best combinations of 2-D structure prediction and 3-D structure prediction programs. Subsequently, the candidate crRNA structures were docked in silico with the corresponding Cas13 protein using HDOCK, which showed the best performance in the validation study. The template-free parameter was used and the 10 best models were retained from the in silico experiments for evaluation.

Since the candidate dataset has no experimentally validated structures, the in silico docking experiments of the candidate crRNA structures were evaluated by comparing with the GT crRNA-Cas13 structures as well as by visual analysis of the human experts. First, we applied K-means clustering of the 10 best models obtained from the in silico docking experiments of each candidate crRNA structure [48]. We used the center of mass of each crRNA as a representative position of the crRNA in the 3-D space. The Euclidean distance of each candidate crRNA model to the GT crRNA model was calculated, and the number of clusters was determined in proportion to the number of candidate crRNAs for each Cas13 protein, which was 20, 80, and 10 for Cas13a, Cas13b, and Cas13d, respectively. From the distance calculations, the closest cluster to the GT crRNA was found for each Cas13 protein, and multiple sequence alignments (MSAs) were performed on the closest cluster for each docking result. Next, the individual candidate crRNAs with the closest docking position to the GT crRNA were found, with the number of individuals also determined proportionally to the size of the candidate dataset for each protein. Followingly, the human experts used the 3-D visualization to validate the candidate crRNA docking results in comparison to the GT results.

## Results

### RNA secondary structures of CRISPR repeats differ by the prediction program

The predicted 2-D structures of the crRNAs in the validation dataset were saved in the dot-bracket format, which is used by every RNA 2-D program (Table S8). None of the crRNAs were predicted to have the same 2-D structure by all eight prediction programs. For most cases, the number of different 2-D structures predicted per CRISPR repeat sequence was between three different structures (5WLH, 6VRB, 6IV8_B, 6IV8_D, 6IV9) and four different structures (5W1H, 5W1I_B, 5W1I_D, 7OS0_D, 7OS0_I, 6DTD, 6E9E). Some crRNAs had more variability in the predicted 2-D structures. For the crRNA of 6AAY, only CentroidFold and CONTRAfold predicted the same RNA 2-D structure. RNAfold and MXfold2 also formed a separate group, as well as RNAstructure and RNAshapes. IPknot and ContextFold each predicted a unique 2-D structure that was different from all the other predictions. Consequently, five different 2-D structures were predicted for this crRNA. For the other crRNAs, the predicted 2-D structures had less variability. For the crRNAs of 5WTK, 5XWY and 6VRC, all the programs predicted the same 2-D structure for each CRISPR repeat sequence except ContextFold.

In overall, ContextFold and CONTRAfold mostly predicted 2-D structures that were more complex with a higher number of paired bases than the rest of the programs. RNAfold, RNAstructure, MXfold2 and RNAshapes predicted the same 2-D structure in most cases. It was also notable that the complexity of the 2-D structures tended to increase from the crRNAs associated with Cas13a, Cas13d to Cas13b. However, there were no consistent patterns to how the eight programs predicted the crRNAs; thus it is important to evaluate the 2-D structure prediction from multiple programs when predicting the structure of an RNA sequence.

### RNA tertiary structures of CRISPR repeats are highly dependent on the predicted secondary structures

The predicted crRNA 3-D structures from RNAComposer and Rosetta were superimposed with the corresponding GT structure using the evaluation programs (PyMOL align, PyMOL super and SETTER) to obtain the RMSD values. For each superimposition of two 3-D structures, the mean and the SD from the three replicate runs were calculated. For SETTER, the RMSD values varied while those of PyMOL align and PyMOL super stayed constant. The RMSD values across all the crRNAs are visualized as a heatmap for each evaluation program (Figures 3, S4-S7). The lower the RMSD value, the smaller the average distance between the atoms of the superimposed structures, and the darker the color in the heatmap.

**Figure 3:**
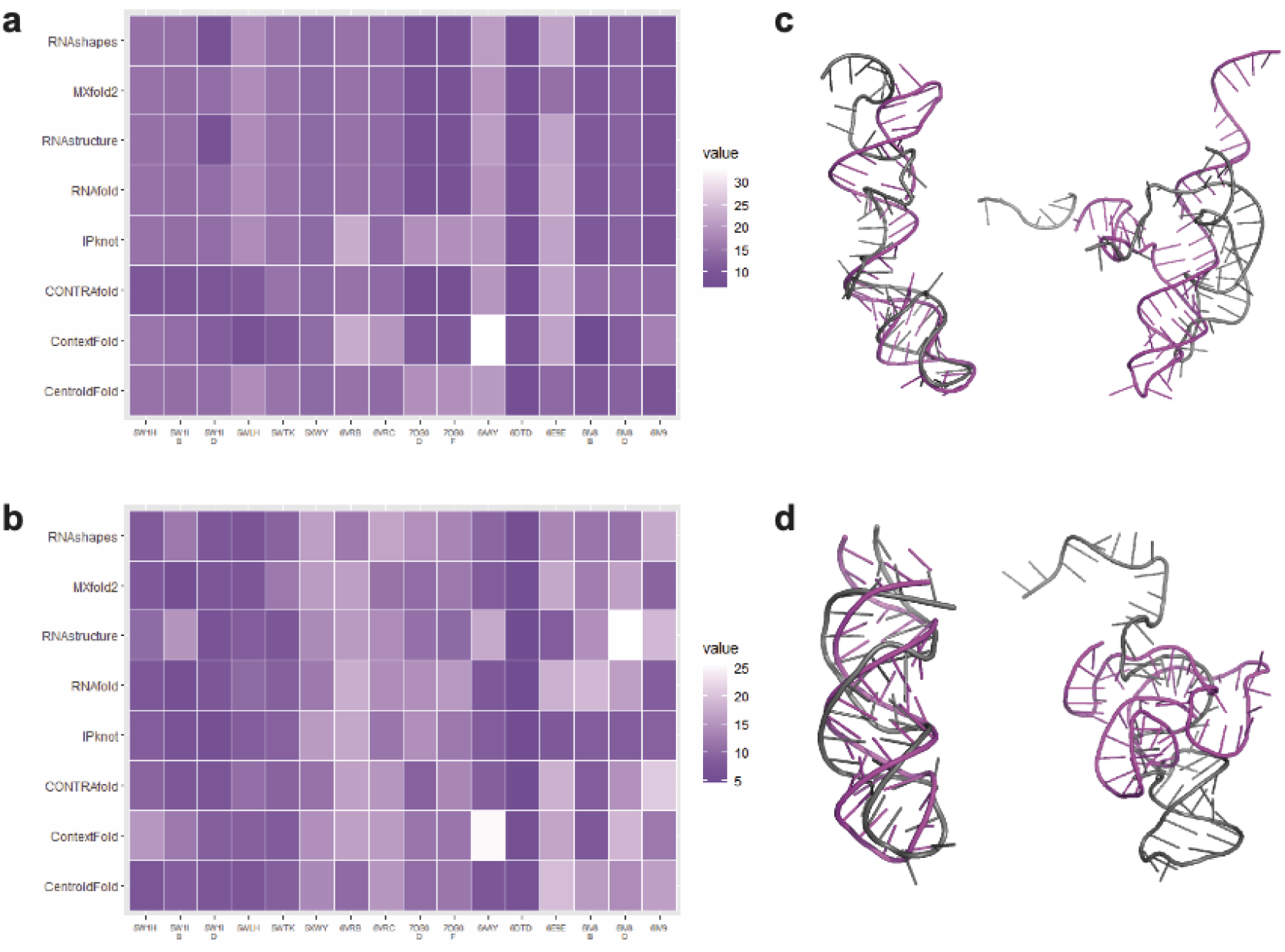
Performance analysis of RNA structure prediction of CRISPR repeats with PyMOL align. Heatmap of the means of the RMSD values by superimposition of each predicted crRNA 3-D structure with the ground truth (GT) structure. The RNA 2-D structure prediction programs are shown on the y-axis, and the PDB name of each Cas13 protein is shown on the x-axis, with the RNA 3-D structure program as **a**. RNAComposer and **b**. Rosetta. **c**. Superimposition of the GT structure 6IV8_B predicted by ContextFold and RNAComposer (best), and 6AAY predicted by ContextFold and RNAComposer (worst). **d**. Superimposition of the GT structure 6DTD predicted by RNAstructure and Rosetta (best), and 6IV8_D predicted by RNAstructure and Rosetta (worst). Grey = GT structure; Magenta = predicted structure

For RNAComposer, the evaluation programs of the superimposition between the predicted 3-D structures and the GT structures varied vastly according to the 2-D structure prediction program used in combination. With the evaluation program of PyMOL align, CONTRAfold gave the lowest RMSD value on average (Table 1), while CentroidFold and IPknot mostly showed lighter shades in the heatmap rows (Figure 3). Interestingly, ContextFold gave both the highest RMSD value and the lowest RMSD value (6AAY and 6IV8_B, respectively). Regarding the performance consistency, ContextFold had the highest SD value. Overall, MXfold2 had the lowest SD value and the second lowest average RMSD value. Thus, the evaluation program of PyMOL align assessed MXfold2 to be the best 2-D structure prediction program in combination with RNAComposer. From the evaluations with PyMOL super and SETTER, the best 2-D prediction programs in combination with RNAComposer were found to be ContextFold and CONTRAfold, respectively (Tables S9-S10).

**Table 1.** Superimposition performance of the predicted crRNA structure with the ground truth crRNA structure. Performance of the RNA 2-D structure prediction programs in combination with RNAComposer or Rosetta when superimposed to the GT 3-D structure of the experimentally validated crRNAs. The mean and the standard deviation of the root-mean-square deviation (RMSD) values were calculated from three replicate runs of PyMOL align across all the predicted structures.

| RNA 3-D structure prediction program | RNA 2-D structure prediction program | Mean RMSD value (Å) | Standard deviation of RMSD values (Å) |
| --- | --- | --- | --- |
| RNAComposer | CentroidFold | 14.5751 | 3.4605 |
|  | ContextFold | 15.0601 | 6.7700 |
|  | CONTRAFold | 13.0886 | 3.9715 |
|  | IPknot | 15.5927 | 4.3685 |
|  | RNAfold | 13.8130 | 4.0411 |
|  | RNAstructure | 13.5431 | 4.2974 |
|  | MXfold2 | 13.3285 | 3.3359 |
|  | RNAshapes | 13.4816 | 4.3896 |
| Rosetta | CentroidFold | 11.4316 | 3.9693 |
|  | ContextFold | 13.2483 | 4.9258 |
|  | CONTRAFold | 11.8012 | 4.3517 |
|  | IPknot | 9.9731 | 3.4309 |
|  | RNAfold | 11.9661 | 4.6712 |
|  | RNAstructure | 12.8113 | 5.3425 |
|  | MXfold2 | 11.1416 | 3.8934 |
|  | RNAshapes | 11.8083 | 3.5449 |

For Rosetta, the performance of the 3-D structure prediction program also varied vastly according to the 2-D structure prediction program used in combination. With the evaluation program of PyMOL align, IPknot gave the lowest RMSD value on average (Table 1), while ContextFold and RNAstructure mostly showed lighter shades in the heatmap rows (Figure 3). Interestingly, RNAstructure both gave the highest RMSD value and the lowest RMSD value (6IV8_B and 6DTD, respectively). Regarding the performance consistency, RNAstructure had the highest SD value. Overall, IPknot had the lowest SD value and also the lowest average RMSD value. Thus, the evaluation program of PyMOL align assessed IPknot to be the best 2-D structure prediction program in combination with Rosetta. From the evaluations with PyMOL super and SETTER, the best 2-D program in combination with Rosetta was found to be RNAfold in both cases (Tables S11-S12).

### crRNA structures of the validation dataset confirm in silico docking with Cas13 proteins

The in silico docking experiments of the validation dataset using HADDOCK showed low performance when the docked model was compared to the GT structure with the CAPRI docking scores (Figure S8). PyDockDNA showed better performance by correctly docking one receptor-ligand pair for every 3-5 pairs. Surprisingly, HDOCK was able to dock all the pairs with high accuracy, regardless of the random rotations. In terms of the iRMSD values, HADDOCK, PyDockDNA and HDOCK achieved on average 28.05, 12.73, and 0.11, with the standard deviations of 5.03, 7.30, and 0.05, respectively, across all the crRNAs in the validation dataset. From the validation study, HDOCK was found to be the best performing in silico docking software for the Cas13 proteins and crRNAs. Due to the exceptional performance, further in silico docking experiments were conducted only with HDOCK.

Next, we used the predicted 3-D structures of the crRNAs in the validation dataset to perform in silico docking with the Cas13a proteins and Cas13b/d proteins (Figure S9 and S10, respectively). Subsequently, the accuracy of the docked model using the predicted crRNA structures was analyzed using the CAPRI docking scores (Table S13). For all the crRNAs, the docking scores of iRMSD were above the acceptable threshold (4 Å) when the RNA 3-D structures were predicted with different combinations of the 2-D and 3-D structure prediction programs (Figures S9-S10). This result was surprising as the previous in silico docking experiments using the GT crRNA structures had resulted in the near-perfect docking score of iRMS (∼0 Å). However, four crRNAs (5WLH, 5W1H, 5W1I_AB, and 5W1I_CD) gave better docking scores than the others, and it was notable that these crRNAs also had better experimental resolutions (Table S2).

As the quality of the GT crRNA structures depended on the experimental condition, we decided to only keep these four crRNAs with the best experimental resolution in further studies [49]. Another adjustment was to consider the 10 best models generated from the in silico docking experiments rather than retaining the best model only. Additionally, the CRISPR repeat sequences for these crRNAs were shortened to match the visible part of the GT structures before predicting the 3-D structures. In the subsequent experiments, some of the docking scores reached an acceptable threshold in terms of the iRMSD scores (Table 2). From visualization by superimposing with the GT structure, the best model selected by the minimal docking score and the human experts were compared. As shown in Figure 4, the best model from the in silico experiments between these two methods coincided, except for 5W1H. Furthermore, the best model of the crRNA associated with 5WLH superimposed almost perfectly with the GT structure. Finally, we performed the in silico docking experiments with both template-free and template-based settings, which resulted in no significant improvement in the docking performance when the template-based setting was used (Figures 4, S11-S12). Thus, all the subsequent in silico docking experiments with the candidate dataset with HDOCK were conducted template-free.

**Table 2.**
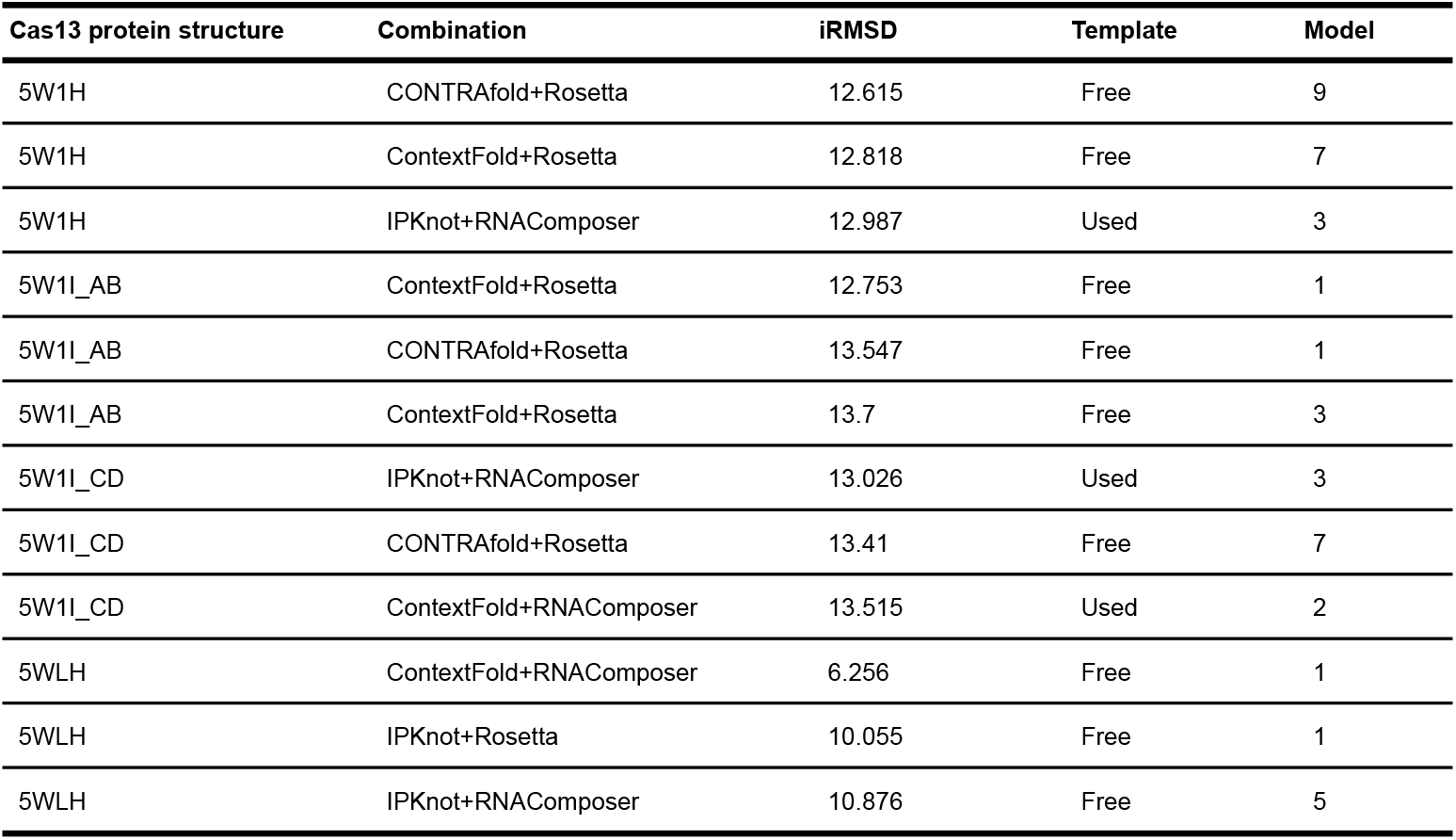
Performance analysis of in silico docking of the predicted crRNA structures with the Cas13a proteins. The three best docking models of the Cas13a proteins with the best experimental resolution were given in terms of iRMSD when superimposed with the GT structures (5W1H, 5W1I_AB, 5W1I_CD, 5WLH). The Model column refers to the *n*^*th*^ best model given by HDOCK in each in silico docking experiment.

| Cas13 protein structure | Combination | iRMSD | Template | Model |
| --- | --- | --- | --- | --- |
| 5W1H | CONTRAFold+Rosetta | 12.615 | Free | 9 |
| 5W1H | ContextFold+Rosetta | 12.818 | Free | 7 |
| 5W1H | IPKnot+RNAComposer | 12.987 | Used | 3 |
| 5W1I_AB | ContextFold+Rosetta | 12.753 | Free | 1 |
| 5W1I_AB | CONTRAFold+Rosetta | 13.547 | Free | 1 |
| 5W1I_AB | ContextFold+Rosetta | 13.7 | Free | 3 |
| 5W1I_CD | IPKnot+RNAComposer | 13.026 | Used | 3 |
| 5W1I_CD | CONTRAFold+Rosetta | 13.41 | Free | 7 |
| 5W1I_CD | ContextFold+RNAComposer | 13.515 | Used | 2 |
| 5WLH | ContextFold+RNAComposer | 6.256 | Free | 1 |
| 5WLH | IPKnot+Rosetta | 10.055 | Free | 1 |
| 5WLH | IPKnot+RNAComposer | 10.876 | Free | 5 |

**Figure 4:**
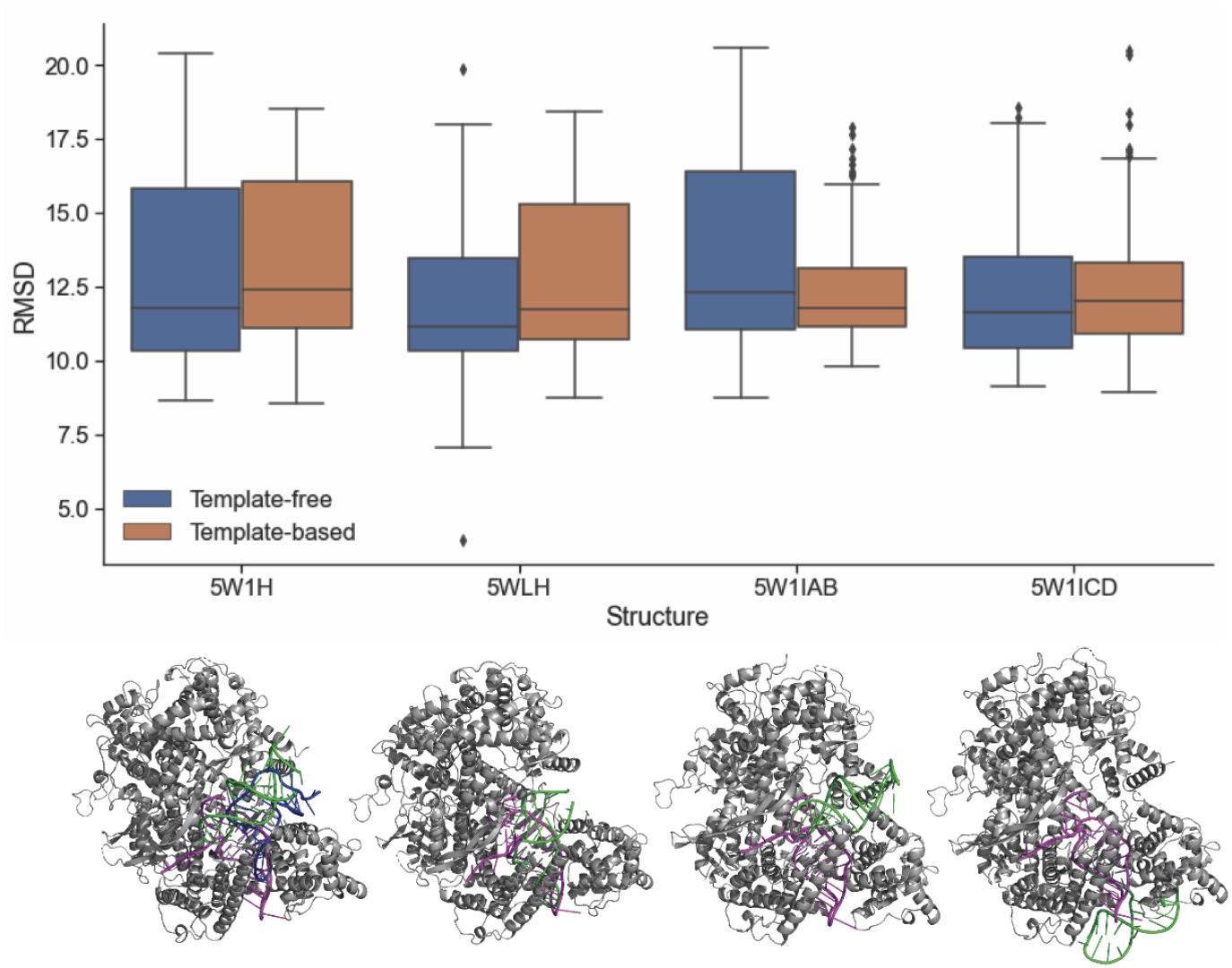
In silico docking evaluation of the Cas13a proteins of the best experimental resolution. The iRMSD from the in silico docking experiments of the crRNAs with the Cas13a protein using HDOCK. The 10 best models were retained from HDOCK and the experiments were performed template-free or template-based. Each box represents the results of 60 docking experiments. The 3-D structure below each box shows the GT structure (magenta), the computer selected best model (blue), and the human selected best model (green) docked on the corresponding Cas13a protein (grey). Except for 5W1H, the computer selected best model coincided with the human selected best model (green).

### Some candidate crRNAs had optimal in silico docking with Cas13 proteins

The predicted 3-D structures of the candidate crRNAs were docked in silico with the corresponding Cas13 protein using the optimized pipeline. As the 10 best models were retained for each docking experiment, there were many docked models to be evaluated by the human experts in the absence of the GT structures. Thus, we first summarized each docking result as the centre of mass in the spatial coordinates calculated from all atoms of the docked macromolecular structure model (Figure 5). This step enabled a visual summary of the in silico docking experiments, and calculation of the closest clusters or individuals to the GT crRNA in the spatial coordinates. As the coordinates of the Cas13 protein remained constant, the docked models of each candidate crRNAs near the GT crRNA had the potential to interact optimally with the binding domains. For example, the N-terminal domain (NTD) and the Helical-1 domain were previously found to form the crRNA-recognition (REC) lobe of Cas13a [49], and we only considered the candidate crRNAs which docked near these domains by selecting the closest clusters or individuals to the GT crRNA. We ranked the best candidate crRNAs for each Cas13 protein from the candidate dataset by calculating the Euclidean distance of each docked model to the GT, which determined the best cluster of the candidate crRNAs (containing 10-50 docked models for each Cas13) and the best individual candidate crRNAs (containing 10-50 docked models for each Cas13). The average distance between each GT crRNA and the closest cluster was 17.15 ± 7.09 Å, and the average distance between each GT crRNA and the closest individual candidate crRNAs was 16.69 ± 5.26 Å. Using a two-sided Student’s t-test, we found that the difference between the average individual and the average cluster distances was not significant (p=0.54, Table S17).

**Figure 5:**
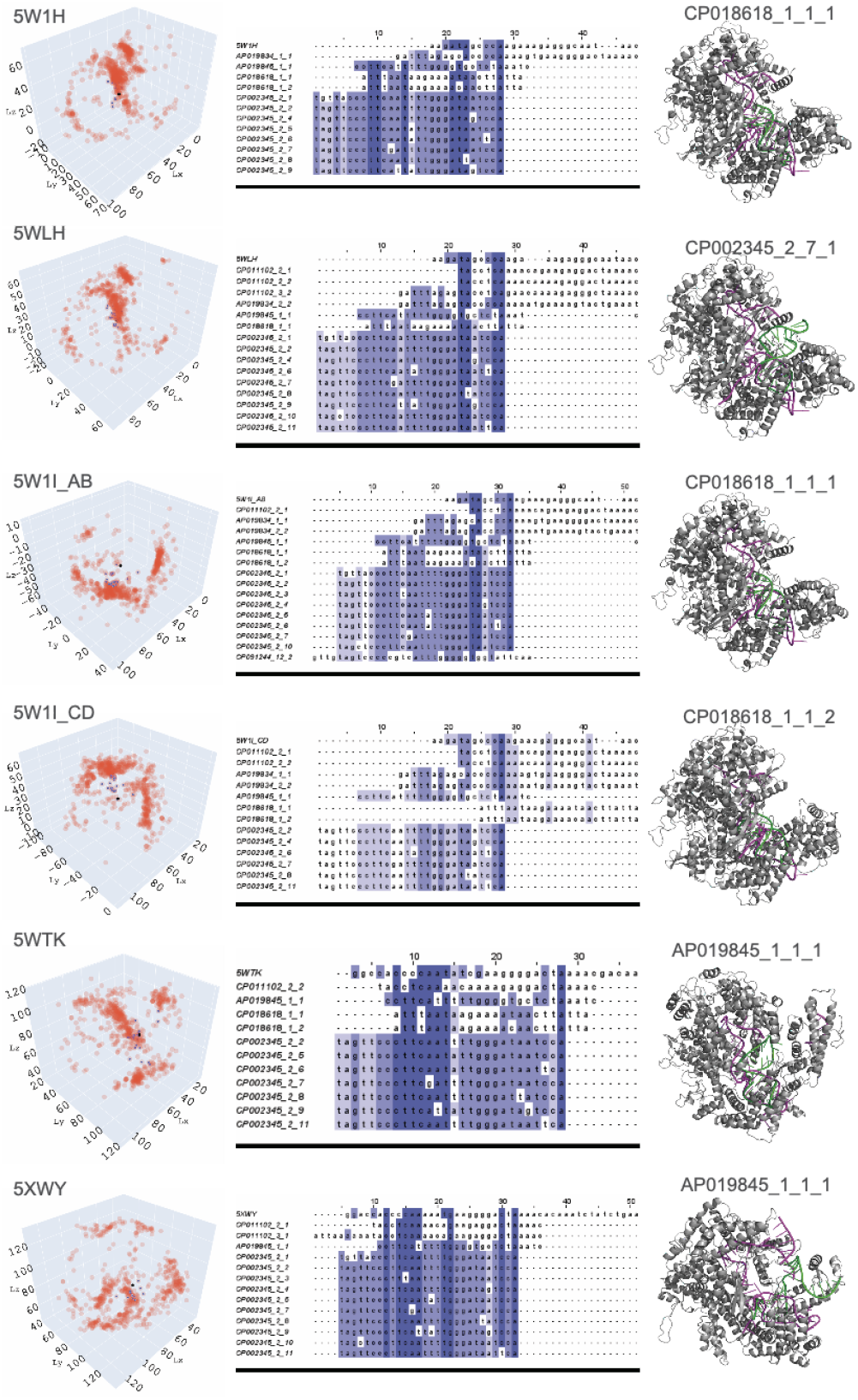
Best crRNA candidates for Cas13a protein from in silico docking experiments. The 3-D visualization shows the spatial coordinates of each crRNA candidate model (only 10 best models were taken into account) after in silico docking with the corresponding Cas13 protein. Each pink dot represents the centre of mass calculated from all atoms of the macromolecular structure model for each crRNA candidate. The GT crRNA is marked as a black dot, and its closest crRNA candidates in terms of Euclidian distance are highlighted as blue dots. The multiple sequence alignment compares the RNA sequences of the GT crRNA with those of the best crRNA candidates. The different shades of blue show the percentage identity, with the identity threshold set to 50%, highlighting variations in the RNA sequences. The 3-D structure shows an example of docking between the receptor (Cas protein) and the ligand (crRNA). The Cas proteins are in grey, the GT crRNA is colored in magenta, and the best crRNA candidate model is highlighted in green with its identifier given above (GenomeID_CRISPRarray_CRISPRrepeat_modelnumber).

The docking results of the selected candidate crRNAs were further evaluated through visual inspection by the human experts (Figures S13-15). From the visualization analysis, the docking results of the best individual crRNAs were found to show better docking than those of the best cluster crRNAs, in terms of both position and direction. Among the 321 docked pairs of the closest clusters, only 8 of them received the best docking score from the human experts, while 77 from the 370 closest individuals received the best docking score (Table S14-S16). It is notable that not all candidate crRNAs with the best docking result came from the best model generated by the in silico docking software. For example, the 8th best model of the candidate crRNA of CP002345_2_11 was evaluated to dock very well with the corresponding Cas protein of 5WTK (Table 3). This result shows the importance of retaining at least the 10 best models from the in silico docking experiments to be analyzed further by the human experts, to ensure that the best docking model of the ligand reflects the biological information on the receptor.

**Table 3.** Best candidate crRNAs from in silico docking with Cas13 proteins. A list of crRNA candidates for each Cas13a protein with optimal in silico docking, selected using the optimized pipeline and calculating the distance to the GT crRNA. The Model column refers to the *n*^*th*^ best model given by HDOCK in each in silico docking experiment. The Distance column shows the Euclidean distance of each docked model to the GT crRNA. The Human expertise column indicates the following 3-D visual assessment by the human expert: 0 = docked in the same region and in a similar direction as GT 1 = partially docked in the similar region and in a similar direction as GT

| Best model | Cas13 protein structure | Candidate crRNA | Model | Distance (Å) | Cluster | Human expertise |
| --- | --- | --- | --- | --- | --- | --- |
| RNAC_5W1HvsAP019845_1_1_1 | 5W1H | AP019845_1_1 | 1 | 15.42061 | 8 | 1 |
| RNAC_5W1HvsAP019845_1_1_5 | 5W1H | AP019845_1_1 | 5 | 15.91598 | 6 | 1 |
| RNAC_5W1HvsCP002345_2_4_1 | 5W1H | CP002345_2_4 | 1 | 18.63954 | 16 | 1 |
| RNAC_5W1HvsCP002345_2_9_10 | 5W1H | CP002345_2_9 | 10 | 8.592774 | 3 | 0 |
| Rose_5W1HvsAP019834_1_1_1 | 5W1H | AP019834_1_1 | 1 | 17.97766 | 3 | 1 |
| Rose_5W1HvsAP019845_1_1_3 | 5W1H | AP019845_1_1 | 3 | 8.767469 | 16 | 0 |
| Rose_5W1HvsAP019845_1_1_7 | 5W1H | AP019845_1_1 | 7 | 16.31799 | 3 | 1 |
| Rose_5W1HvsCP002345_2_6_1 | 5W1H | CP002345_2_6 | 1 | 10.5807 | 3 | 0 |
| Rose_5W1HvsCP002345_2_7_1 | 5W1H | CP002345_2_7 | 1 | 10.28279 | 3 | 0 |
| Rose_5W1HvsCP018618_1_1_1 | 5W1H | CP018618_1_1 | 1 | 11.36411 | 3 | 0 |
| RNAC_5W1I_ABvsAP019834_2_2_1 | 5W1I_AB | AP019834_2_2 | 1 | 15.41219 | 1 | 1 |
| RNAC_5W1I_ABvsAP019845_1_1_1 | 5W1I_AB | AP019845_1_1 | 1 | 15.4393 | 7 | 1 |
| RNAC_5W1I_ABvsAP019845_1_1_2 | 5W1I_AB | AP019845_1_1 | 2 | 16.81572 | 15 | 1 |
| RNAC_5W1I_ABvsCP091244_12_2_8 | 5W1I_AB | CP091244_12_2 | 8 | 19.48762 | 9 | 1 |
| Rose_5W1I_ABvsAP019834_1_1_7. | 5W1I_AB | AP019834_1_1 | 7 | 18.04252 | 9 | 1 |
| Rose_5W1I_ABvsAP019845_1_1_2 | 5W1I_AB | AP019845_1_1 | 2 | 17.05391 | 9 | 1 |
| Rose_5W1I_ABvsCP002345_2_5_4 | 5W1I_AB | CP002345_2_5 | 4 | 14.53866 | 1 | 1 |
| Rose_5W1I_ABvsCP002345_2_6_2 | 5W1I_AB | CP002345_2_6 | 2 | 10.42858 | 1 | 0 |
| Rose_5W1I_ABvsCP002345_2_6_5 | 5W1I_AB | CP002345_2_6 | 5 | 13.87067 | 1 | 0 |
| Rose_5W1I_ABvsCP002345_2_7_2 | 5W1I_AB | CP002345_2_7 | 2 | 10.77602 | 1 | 0 |
| Rose_5W1I_ABvsCP018618_1_1_1 | 5W1I_AB | CP018618_1_1 | 1 | 10.17277 | 1 | 0 |
| RNAC_5W1I_CDvsAP019834_2_2_3 | 5W1I_CD | AP019834_2_2 | 3 | 14.82603 | 18 | 0 |
| RNAC_5W1I_CDvsAP019845_1_1_1 | 5W1I_CD | AP019845_1_1 | 1 | 15.82544 | 19 | 1 |
| RNAC_5W1I_CDvsAP019845_1_1_10 | 5W1I_CD | AP019845_1_1 | 10 | 11.31408 | 18 | 1 |
| Rose_5W1I_CDvsAP019834_1_1_5 | 5W1I_CD | AP019834_1_1 | 5 | 18.23438 | 2 | 1 |
| Rose_5W1I_CDvsAP019845_1_1_9 | 5W1I_CD | AP019845_1_1 | 9 | 18.28236 | 2 | 1 |
| Rose_5W1I_CDvsCP002345_2_4_4 | 5W1I_CD | CP002345_2_4 | 4 | 10.64548 | 18 | 0 |
| Rose_5W1I_CDvsCP002345_2_6_1 | 5W1I_CD | CP002345_2_6 | 1 | 10.06707 | 18 | 0 |
| Rose_5W1I_CDvsCP002345_2_6_6 | 5W1I_CD | CP002345_2_6 | 6 | 13.18188 | 18 | 0 |
| Rose_5W1I_CDvsCP002345_2_6_7 | 5W1I_CD | CP002345_2_6 | 7 | 10.06989 | 18 | 0 |
| Rose_5W1I_CDvsCP002345_2_7_1 | 5W1I_CD | CP002345_2_7 | 1 | 10.96614 | 18 | 0 |
| Rose_5W1I_CDvsCP011102_2_1_3 | 5W1I_CD | CP011102_2_1 | 3 | 19.27656 | 18 | 1 |
| Rose_5W1I_CDvsCP018618_1_1_2 | 5W1I_CD | CP018618_1_1 | 2 | 8.83881 | 18 | 0 |
| RNAC_5WLHvsAP019834_2_2_2 | 5WLH | AP019834_2_2 | 2 | 15.66461 | 2 | 1 |
| RNAC_5WLHvsAP019845_1_1_1 | 5WLH | AP019845_1_1 | 1 | 13.99831 | 10 | 1 |
| RNAC_5WLHvsCP002345_2_1_4 | 5WLH | CP002345_2_1 | 4 | 14.43221 | 2 | 1 |
| RNAC_5WLHvsCP011102_2_2_4 | 5WLH | CP011102_2_2 | 4 | 13.4716 | 2 | 0 |
| Rose_5WLHvsAP019845_1_1_1 | 5WLH | AP019845_1_1 | 1 | 16.64761 | 2 | 1 |
| Rose_5WLHvsAP019845_1_1_3 | 5WLH | AP019845_1_1 | 3 | 12.08608 | 2 | 1 |
| Rose_5WLHvsAP019845_1_1_4 | 5WLH | AP019845_1_1 | 4 | 14.27125 | 2 | 1 |
| Rose_5WLHvsCP002345_2_6_1 | 5WLH | CP002345_2_6 | 1 | 11.24522 | 2 | 0 |
| Rose_5WLHvsCP002345_2_6_3 | 5WLH | CP002345_2_6 | 3 | 14.17092 | 2 | 0 |
| Rose_5WLHvsCP002345_2_7_1 | 5WLH | CP002345_2_7 | 1 | 11.88083 | 2 | 0 |
| Rose_5WLHvsCP011102_3_2_5 | 5WLH | CP011102_3_2 | 5 | 16.4966 | 2 | 1 |
| RNAC_5WTKvsAP019845_1_1_1 | 5WTK | AP019845_1_1 | 1 | 23.34363 | 9 | 1 |
| Rose_5WTKvsAP019845_1_1_1 | 5WTK | AP019845_1_1 | 1 | 3.64798 | 5 | 0 |
| Rose_5WTKvsAP019845_1_1_2 | 5WTK | AP019845_1_1 | 2 | 1.859212 | 5 | 0 |
| Rose_5WTKvsAP019845_1_1_3 | 5WTK | AP019845_1_1 | 3 | 3.46364 | 5 | 0 |
| Rose_5WTKvsCP002345_2_11_8 | 5WTK | CP002345_2_11 | 8 | 4.103924 | 5 | 0 |
| Rose_5WTKvsCP002345_2_5_1 | 5WTK | CP002345_2_5 | 1 | 9.187617 | 5 | 1 |
| Rose_5WTKvsCP002345_2_6_1 | 5WTK | CP002345_2_6 | 1 | 2.796534 | 5 | 0 |
| Rose_5WTKvsCP002345_2_6_2 | 5WTK | CP002345_2_6 | 2 | 2.4628 | 5 | 0 |
| Rose_5WTKvsCP002345_2_8_1 | 5WTK | CP002345_2_8 | 1 | 5.023773 | 5 | 0 |
| Rose_5WTKvsCP011102_2_2_1 | 5WTK | CP011102_2_2 | 1 | 6.560322 | 5 | 1 |
| Rose_5WTKvsCP018618_1_2_8 | 5WTK | CP018618_1_2 | 8 | 5.777393 | 5 | 1 |
| RNAC_5XWYvsAP019845_1_1_1 | 5XWY | AP019845_1_1 | 1 | 10.34585 | 2 | 0 |
| RNAC_5XWYvsAP019845_1_1_3 | 5XWY | AP019845_1_1 | 3 | 14.16364 | 6 | 1 |
| Rose_5XWYvsCP011102_2_1_10 | 5XWY | CP011102_2_1 | 10 | 17.63066 | 2 | 0 |

The multiple sequence alignment (MSA) results showed that some candidate crRNAs in the closest individuals to the GT crRNA had very different sequences when compared to the sequence of the GT crRNA (Figure 5). When these crRNAs were visualized, they had the in silico docking models as optimal as the GT crRNA without necessarily having similar sequences. For example, the crRNA of CP018618_1_1 fits perfectly into the docking region of 5W1H, but its sequence has almost no similarity to the sequence of the GT crRNA. Therefore, a selected list of candidate crRNAs is provided as novel CRISPR repeats that have the potential to interact as optimally as the GT crRNAs for each Cas13 protein. These candidate crRNAs have been assessed through in silico docking experiments worthwhile to be further validated through in vitro or in vivo experiments (Table 3).

## Discussion

Conducting in silico experiments that accurately predict the results of laborious and expensive laboratory experiments is a long-standing goal of many computational biologists. The recent advances in computational methods such as machine learning in 3-D protein structure prediction [19,50] and in genomics [51,52] bring the possibility of achieving such challenging tasks closer to reality. In this study, a series of in silico docking experiments were conducted to model the RNA-protein interactions between a Cas protein and a CRISPR repeat. We predicted the RNA 3-D structures of these CRISPR repeats to perform in silico docking with the corresponding Cas13 protein, with the first goal of optimizing the in silico docking pipeline and the second goal of generating a list of potential CRISPR repeats that may interact optimally with the Cas13-based antimicrobial tools.

The first part of the study used the validation dataset of the crRNA-Cas complexes with the experimentally validated structures to optimize the computational pipeline of in silico docking, given a CRISPR repeat sequence and a Cas protein structure as the input data. This optimization followed the iterative process of selecting the best combination of the 2-D and 3-D structure prediction programs through evaluating the predicted RNA structure against the GT RNA structure. Subsequently, the GT structures of the crRNA-Cas complexes were utilized to select the best in silico docking software with Cas proteins as the receptor and crRNAs as the ligand. This optimized pipeline was further tested using the predicted 3-D structures of crRNAs to assess the impact of structure prediction on the performance of in silico docking, as compared to the GT crRNA 3-D structures. We quantified the impact of inaccuracy arising from the two-step process of the 2-D and 3-D structure prediction, and optimized the subsequent in silico docking experiments. For this optimization, we achieved satisfactory results according to the evaluation metrics utilized in the CAPRI community-wide experiment that aims at modelling interactions based on the 3-D structure of macromolecules [53].

The second part of this study aimed at finding candidate crRNAs that are predicted to have better or equivalent interactions with the Cas13 proteins as the original crRNAs in the validation dataset. This candidate dataset was generated by curating a set of CRISPR repeat sequences that are colocalized with the Cas13 system, as the previous studies demonstrated that the proximal colocalization of CRISPR arrays is an indication of association to the Cas system [54]. Using the previously optimized pipeline, we conducted in silico docking experiments by first predicting the 3-D structure of these candidate crRNAs, which were subsequently docked with the corresponding Cas13 protein. The in silico docking experiments were evaluated by comparing the spatial coordinates of the docked RNA models with those of the GT crRNA, relative to the corresponding Cas13 protein. This evaluation step enabled the best docked model to be selected efficiently by ranking hundreds of docked models in terms of the closest Euclidean distance to the GT crRNA, thus to the crRNA-recognition domains of the Cas13 protein. The final step of the candidate study was the intervention of human expertise by visualizing these docked models to evaluate the receptor-ligand binding as compared to the interaction of the GT crRNA-protein complexes. Remarkably, we found a number of candidate crRNAs that showed in silico docking comparable to the GT crRNA-protein complexes, despite the dissimilarity in genetic sequence. Given that the 3-D structures of these candidate crRNAs were predicted, this result indicates the potential of these RNA-protein interactions to be more stable than those of the GT crRNA-protein complexes.

The in silico docking experiments with the predicted crRNA structures and the associated Cas13 proteins conducted in this study are significant for the following reasons. With the increasing availability of metagenomic sequencing, CRISPR-Cas systems in nature are discovered to be more diverse, complex and disordered than previously expected, and the associations between CRISPR arrays and Cas proteins are enigmatic in some prokaryotic genomes. Particularly, the recently discovered Cas13 systems that degrade RNAs are found to be different architecture-wise from the previous CRISPR-Cas systems, as indicated by the absence of CRISPR arrays and *cas1*/*cas2* genes in vicinity [26,27]. However, the exact mechanism of how Cas13-based systems can function without the adaptation module and colocalized CRISPR arrays is still an ongoing investigation. Given that the Cas13-based systems have been proposed as alternative antimicrobial tools, it is imperative to investigate the characteristics of these novel CRISPR-Cas systems. This preliminary study is an important step towards designing more effective CRISPR-Cas13-based antimicrobial tools, which may be susceptible to off-targeting events in the presence of crRNAs with higher binding affinity. This problem is even more evident in pathogenic bacteria whose genomes are known to contain several endogenous CRISPR-Cas systems [23]. As future prospects, the selected candidate crRNAs in this study should be tested in vitro and in vivo to validate the outcome of these in silico docking experiments. This experimental validation will elucidate the potential off-target effects of the CRISPR-Cas13-based antimicrobial tools, which would be an important step towards optimizing the crRNA-Cas13 complex to be stable and effective in targeting multidrug-resistant bacteria within the complex environment of human microbiota. Finally, this study reveals a number of aspects of in silico docking that could be improved with further investigations to incorporate recent computational and biological advances. We suggest that predicting receptor-ligand interactions is another biological field where deep-learning applications may become extremely valuable.

## Data Availability

All the input protein sequences are available in CRISPRCasdb (https://crisprcas.i2bc.paris-saclay.fr/MainDb/StrainList). The 3-D structures from the Protein Data Bank (https://www.rcsb.org) were used as ground truth for assessing structure prediction and in silico docking (downloaded on 28/02/2022). Figure 1 was created with BioRender.com. All predicted 3-D structures and docked models are available on our github project page, including 3-D images.

## Code Availability

The codes related to this project are available at our GitHub page (https://github.com/hshimlab/CRISPR-Cas-interaction). 2-D RNA structures were predicted with programs in Table S4 (accessed on 01/March/2022). 3-D RNA structures were predicted with RNAComposer (https://rnacomposer.cs.put.poznan.pl/, accessed on 15/June/2022) and Rosetta (https://www.rosettacommons.org/, accessed on 15/June/2022). In silico docking experiments were modelled with HADDOCK2.4 (https://wenmr.science.uu.nl/5/, accessed on 9/Jan/2022), HDOCK (http://hdock.phys.hust.edu.cn/, accessed on 4/April/2022), PyDockDNA (https://model3dbio.csic.es/pydockdna, accessed on 4/April/2022) and were automated with Python 3.8.3 (https://www.python.org), Selenium v.3.141.0 (https://www.selenium.dev/), ChromeDriver v. 100.0.4896.60 (https://chromedriver.chromium.org/), urllib.request v.3.8 (https://pypi.org/project/urllib3/), PyVirtualDisplay v.3.0 (https://pypi.org/project/PyVirtualDisplay/), pandas v.1.0.5 (https://github.com/pandas-dev/pandas), and PyMOL v.2.4.1 (https://pymol.org/2/). For MSA, we used MAFFT v.7.490 (https://mafft.cbrc.jp/alignment/software/) for calculation and Jalview v.2 (https://www.jalview.org) for visualization. 3-D Structure visualizations were created in PyMOL v.2.5.2 (https://pymol.org) and Py3DMol v.1.7.0 (https://pypi.org/project/py3Dmol) with Jupyter v.1.0.0 (https://jupyter.org). For data analysis, Python .3.6.4 (https://www.python.org), NumPy v.1.17.5 (https://github.com/numpy/numpy), SciPy v.1.1.0 (https://www.scipy.org), seaborn v.0.9.0 (https://github.com/mwaskom/seaborn), Matplotlib v.3.3.4 (https://github.com/matplotlib/matplotlib), pandas v.0.22.0 (https://github.com/pandas-dev/pandas), and DockQ (https://github.com/bjornwallner/DockQ, accessed on 10/May/2022) were used.

## Acknowledgements

The research and development activities described in this study were funded by Ghent University Global Campus (GUGC), Incheon, Korea.

## Competing interests

None

## Supplementary Information

Supplementary Information will be provided upon request.

## Notes

### Competing Interest Statement

The authors have declared no competing interest.

